# What if you are not certain? A common computation underlying action selection, reaction time and confidence judgment

**DOI:** 10.1101/180281

**Authors:** Vassilios Christopoulos, Vince Enachescu, Paul Schrater, Stefan Schaal

## Abstract

From what to wear to a friend’s party, to whether to stay in academia or pursue a career in industry, nearly all of our decisions are accompanied by a degree of confidence that provides an assessment of the expected outcome. Although significant progress has been made in understanding the computations underlying confidence judgment, the preponderance of studies focuses on perceptual decisions, in which individuals sequentially sample noisy information and accumulate it as evidence until a threshold is exceeded. Once a decision is made, they initiate an action to implement the choice. However, we often have to make decisions during ongoing actions in dynamic environments where the value and the availability of the alternative options can change with time and previous actions. The current study aims to decipher the computations underlying confidence judgment in action decisions that are made in a dynamic environment. Using a reaching task in which movements are initiated to multiple potential targets, we show that action selection, reaction time and choice confidence all emerge from a common computation in which parallel prepared actions compete based on the overall desirability of targets and action plans.

## 1 Introduction

On January 15, 2009, the US Airways flight 1549, a domestic flight from La Guardia Aiport in New York City to Seattle/Tacoma, experienced a complete loss of thrust in both engines after encountering a flock of Canada geese. As the aircraft lost altitude, the air traffic control asked the pilot if he could either return to La Guardia or to land at the nearby Teterboro airport. Having less than 5 minutes after the bird strike to land the plane, the pilot rejected both options, because he was not *confident* that he could make any runway. Instead, he safely glided the plane to ditch in the Hudson river. Later investigation showed that the low altitude and the lack of power on both engines would not allow for a successful landing to either airport. This incident describes a ubiquitous situation in which choice confidence - i.e., the subjective belief that a given action is more *desirable* than any alternative - has a key role in guiding behavior, especially in dynamic decisions that are made under pressure and while acting. Although confidence is an essential component in human behavior, only recently have we begun to decipher the computations underlying confidence. However, most of this understanding has been built on a fairly restrictive experimental paradigm involving simple decisions like perceptual judgments [1–5] and value-based decisions [6] in stable environments where actions occur only after a choice is made.

While many of our decisions are solely based on incoming sensory information, we must often select between competing options by integrating information from disparate sources (e.g., altitude of the plane, thrust of the engines, airplane condition, etc) while acting. In the current study, we aim to elucidate the computations underlying choice confidence, modeling confidence as a belief that an action has an overall better set of outcomes (costs and benefits) than alternatives. We designed a “reach-before-you-know” experiment that involved rapid reaches to two potential targets presented simultaneously in both hemifields [7, 8]. Critically, the actual goal location was not disclosed before the movement onset. Dual-target trials were interleaved with single-target trials in which one target was presented either in the left or the right hemifield. By varying the target probability to induce different levels of uncertainty, we tested how goal location uncertainty influences behavior. We found that when both targets had about the same probability of action, individuals delayed making a decision and moved towards an intermediary location, waiting to collect more information before selecting one of the targets - a spatial averaging strategy reported in previous studies [7,9,10]. On the contrary, when one of the targets had higher probability of action, reaches had faster responses and launched closer to the likely target. These findings suggest that target certainty influences both planning and execution of actions in decisions with multiple competing options. Surprisingly, the relationship between approach direction with reaction time was not fully mediated by the target probability. Instead, when people waited longer to initiate an action, reaches were frequently launched towards an intermediary location between the potential goals, regardless of the target probability.

To better understand the relationships between confidence, reaction time and trajectories, we modeled the decision task within a recently proposed computational theory [11,12]. The theory builds on the affordance competition hypothesis, in which multiple actions are formed concurrently and compete over time until one has a sufficient evidence to win the competition [13, 14]. We replace evidence with desirability - a continuously accumulated quantity that integrates all sources of information about the relative value of an action with respect to alternatives. Reaching movements are generated as a mixture of actions weighted by their relative desirability values. In analogy with the normative evidence accumulation models [4, 15, 16], we determine choice confidence through the desirability values. Ambiguous desirabilities indicate that the net evidence supporting one option over the others is weak and therefore the confidence level about the current best action is low. On the contrary, when one action outperforms the alternatives, the net evidence is strong and choice confidence in high. Therefore, the “winning” action determines the selected target and the reaction time, whereas the “losing” action contributes to the computation of confidence - i.e., the closer the desirability of the non-selected action to the desirability of the selected one, the lower the choice confidence. Because desirability is time- and state-dependent, and action competition often does not end after movement onset, selected actions can be changed or corrected in-flight (i.e., change of mind) when confidence is sufficiently low, and/or in the presence of new incoming information. Hence, the model predicts that both movement direction and reaction time can be used as an easy-to-measure proxies for choice confidence. When people are uncertain about the current best option, decisions are delayed by both moving towards an intermediary location and by having longer reaction time. In contrast, when they are certain, reaches are initiated faster and move directly to a target. Importantly, the model predicts that the association between approach direction and reaction time is not fully mediated by the target certainty. Instead, action competition can diminish choice confidence leading to slower responses regardless of target probability. Overall, model predictions are consistent with human findings providing direct evidence that action selection, reaction time and choice confidence emerge through a common mechanism of desirability-driven competition between parallel prepared actions.

## 2 Results

### 2.1 Behavioral paradigm

A schematic representation of the experimental setup is shown in Fig. 1. Participants were instructed to perform rapid reaches using a robotic manipulandum under a “reach-before-you-know” paradigm [7, 8] in which either one (single-target trials) or two (dual-target trials) potential targets presented simultaneously in opposite hemifields. For dual-target trials, the cues appeared symmetric around the vertical axis of the screen. By varying the number of potential targets and their probabilities, we induce different level of uncertainty to study the computations underlying choice confidence in action decisions. Each participant ran two separate sessions. In the **equiprobable** session, a trial started with participants fixating on a central cross, followed by the presentation of one or two unfilled blue circles in the screen Fig. 2A. When the fixation cue was extinguished, an auditory cue signaled the individuals to initiate their responses. Once the reaching movement exceeded a threshold, one of the targets filled-in black indicating the actual goal location. The **unequiprobable** session was similar to equiprobable except for the dual-target trials, in which one of the potential targets was always assigned with higher probability (0.8) than alternative one (0.2). The targets with the high and low probabilities were indicated by unfilled green and red cues, respectively. In single-target trials (i.e., target probability 1) which were randomly interleaved with the dual-target trials in both sessions, a single unfilled blue cue was presented in the left or the right hemifield. The set of target configurations is shown in Fig. 2B. Participants achieved an overall success rate around 93% and their performance was similar between the two sessions (93% and 90% respectively).

**Figure 1.**
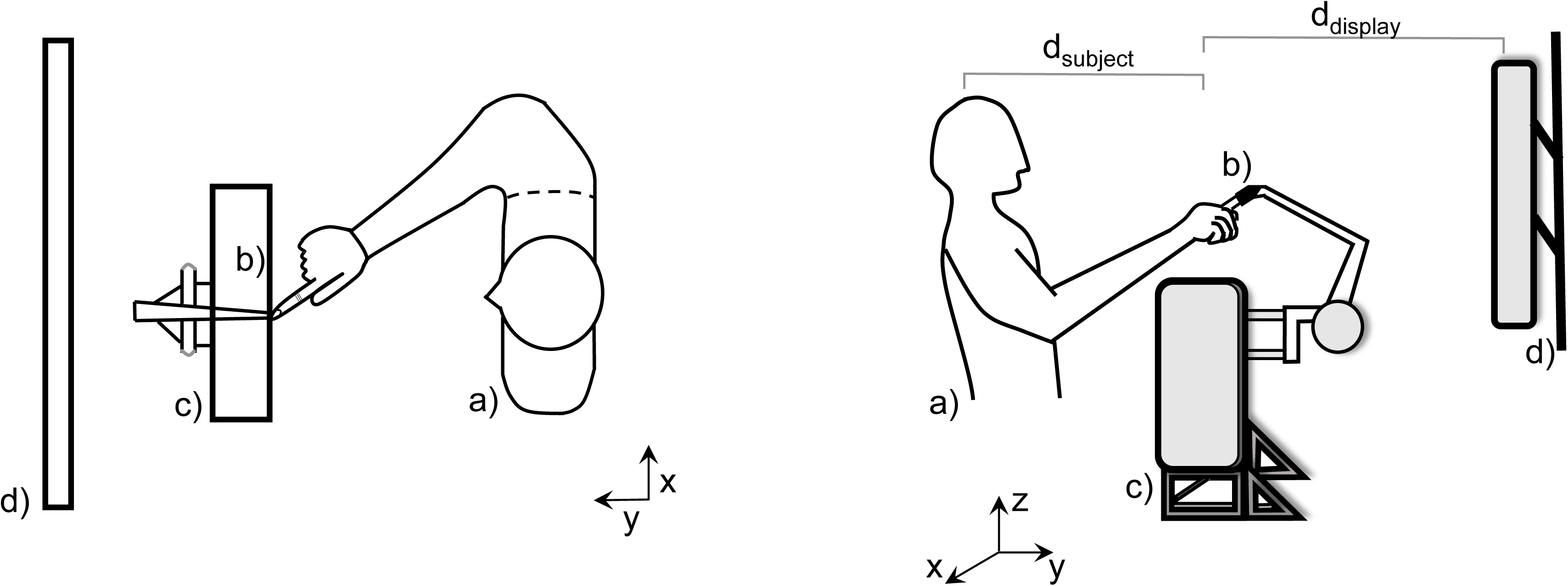
A graphical representation of the experimental setup from two perspectives. Participants (a) were seated directly in front of a Phantom haptic robot (c), with their index finger inserted in a finger-tip adaptor (b) and their midline aligned with the center of an LCD monitor (d). Reaching movements took place in the *x - y* plane, +*y* being towards the screen and +*x* being towards the right hand side of the screen. The distance from the head of the individuals to the finger starting position along the *y* axis was about *d*_*subject*_ = 0.30 m and slightly varied across participants. The distance from the finger starting position to the screen display was *d*_*display*_ = 0.35 m.

**Figure 2.**
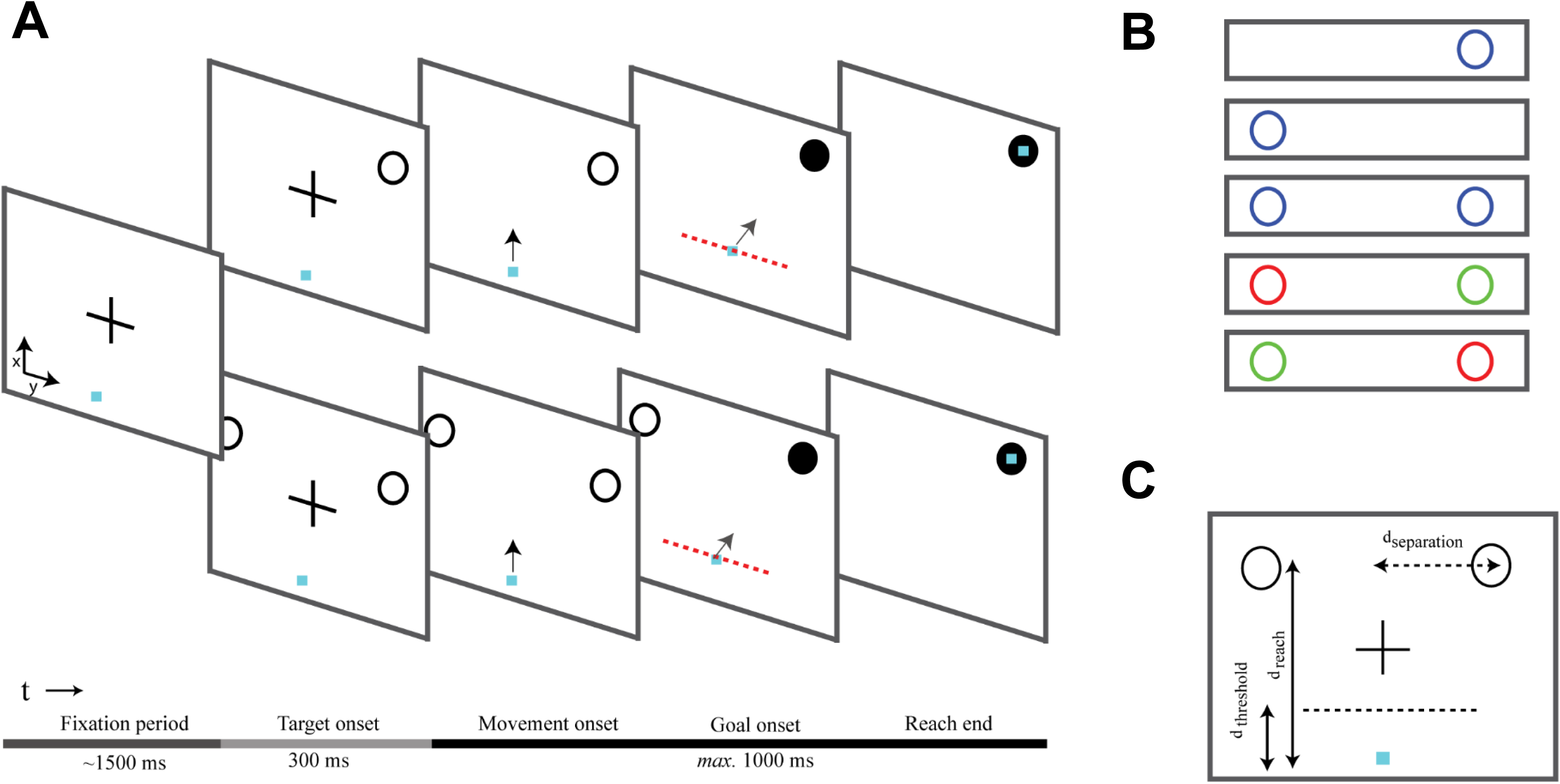
Task design and experimental paradigm. (**A**): A reaching trial started with a fixation cross presented on the center of the screen for about 1.5 s. Then, either a single or two unfilled cues were presented simultaneously in both visual fields. After 300 ms the central fixation cross was extinguished (“go-signal”), and the participants had to perform a rapid reaching movement towards the target(s) within 1 s. Once the reach trajectory crossed a trigger threshold (red discontinuous line), one of the cues (or the single cue) was filled-in black indicating the actual goal location. Responses before the go-signal or reaches that exceeded the maximum movement time (1*s*) were aborted and not used for further analysis. (**B**): The color of the cues in the dual-target trials indicated the target probabilities - blue cues corresponded to equiprobable targets, wheres green and red cues corresponded to targets with 80% and 20% probability, respectively. Single cues always had blue color. (**C**): The distance between the origin and the midpoint of the two cues was *d*_*reach*_ = 0.2 *m*. The distance between the cue and the midpoint was *d*_*separation*_ = 0.15 *m*. The trigger threshold - i.e., distance between the origin and the location that the actual goal location was revealed - was set to *d*_*threshold*_ = 0.05 *m*.

### 2.2 Initial approach direction varies with target probability

Goal location uncertainty is well known to have a strong effect on reach trajectories, where the initial movement trajectory is aimed between targets. This motor behavior, which has been extensively reported before [7, 9, 10, 17], indicates that the approach direction of the initial reaches varies with the target probability, a finding we replicated. Average reach trajectories from a representative participant are illustrated in Figs. 3A and B, when the actual goal was located in the left and the right hemifield, respectively. When there was no uncertainty, reaches were made directly to the goal target (black traces). However, when the goal location was unknown at movement onset, but both targets had the same probability, reaches were aimed to an intermediary position between the potential goal locations (blue traces). These spatially averaged movements were reliably biased towards the side of space with the most likely target (green traces). We measured the approach direction across participants, number of targets and probabilities and found that it is directly correlated with the target certainty (best fit linear regression model; R-square = 0.971, p-value = 0.00212 of the linear coefficient) Fig. 4A. However, we also found that uncertainty has a big impact on reach *initiation*. When people are uncertain about the current best action, they both delayed their decision and moved towards an intermediary location between the targets, a strategy consistent with increasing chances of collecting more information before making a choice.

**Figure 3.**
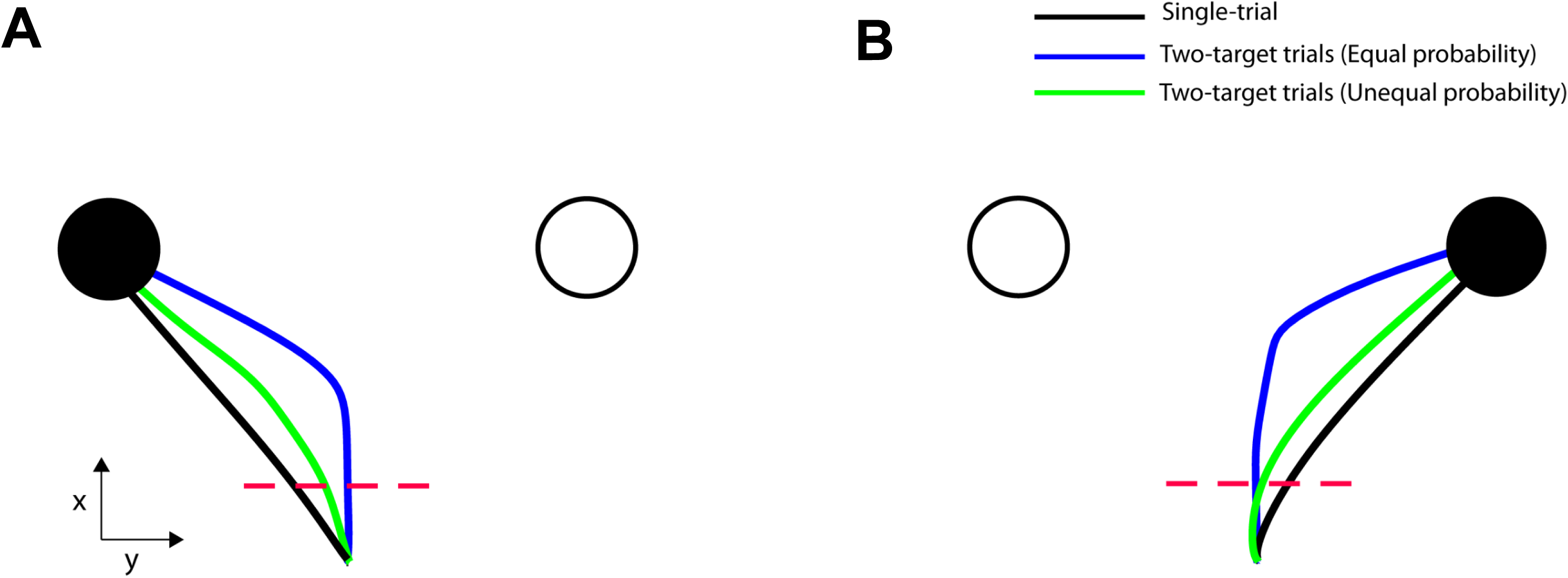
Average reach trajectories from a representative participant. (**A**): Reach trajectories from single-(black trace) and two-target trials with equal (blue trace) and unequal (green trace) probability, with actual goal located in the left hemifield. (**B**): Similar to A but for actual goal located in the right hemifield. Target probability influences the reach trajectories. When people were certain about the goal location, reaches were aimed directly to the target. When they were uncertain, reaches were launched to an intermediary location and then corrected in-flight to the cued target location. The spatially averaged behavior was biased towards the likely target.

### 2.3 Reaction time varies with the target probability

The dual effects of goal uncertainty on reach trajectory and reach initiation timing suggest target certainty is incorporated into both acting (trajectory generation) and planning processes. Intuitively, it is reasonable that target probability influences action planning to delay initiating action when uncertain about the best option. This predicts reaction times (RT) would be a direct function of target probability. On average, this prediction is validated as illustrated in Fig. 4B for single-target trials, two-target trials with equal probability and two-target trials with unequal probability, with RT averaged across participants. While RT is significantly correlated with the target certainty (best fit quadratic regression model; R-square = 0.994, p-value = 0.002 of the quadratic coefficient), a trial-by-trial analysis showed that the effect on initiation timing was indirect and actually mediated by a latent variable influencing both RT and the approach direction of a trajectory.

**Figure 4.**
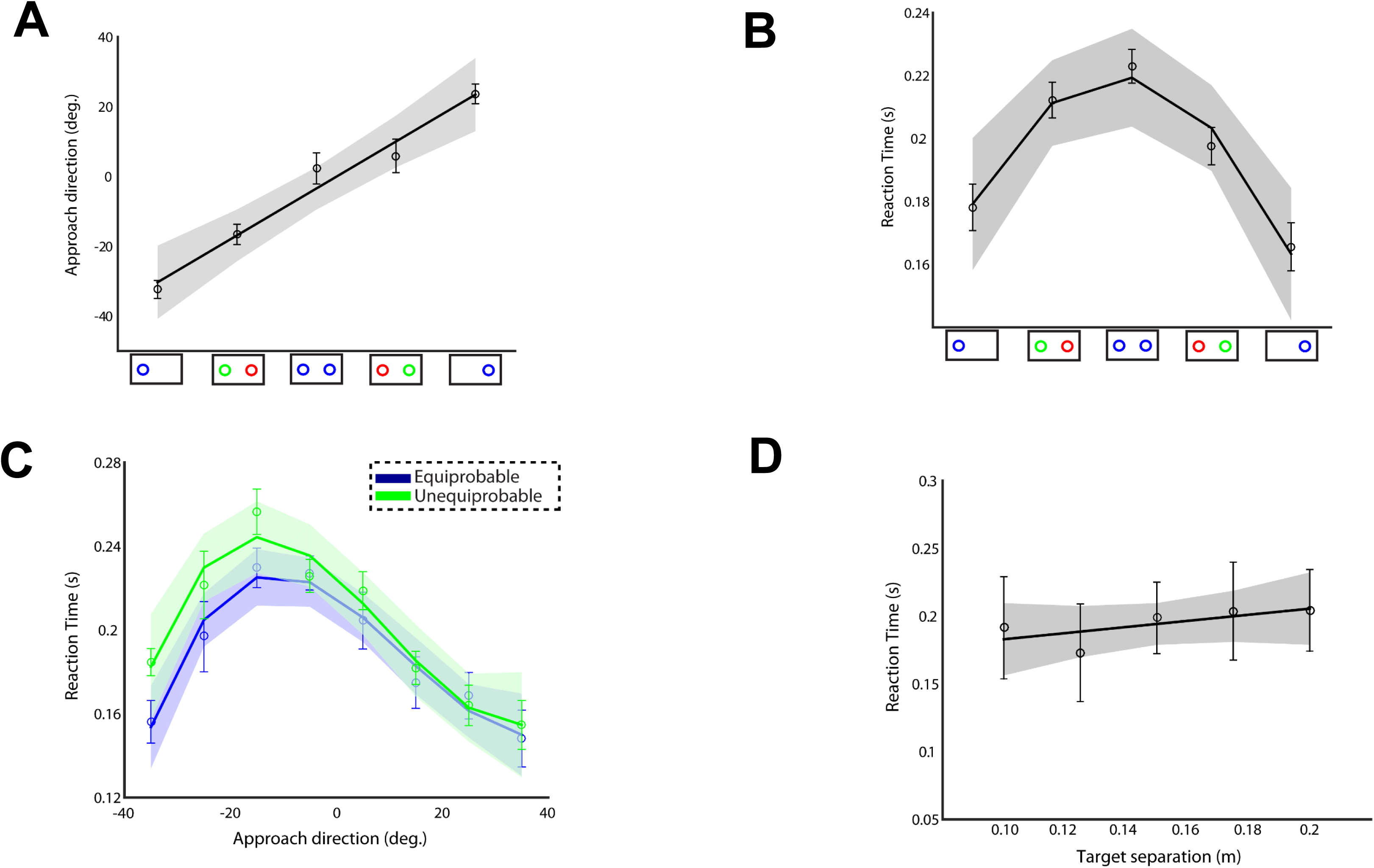
Approach direction and reaction time. (**A**): Approach direction and (**B**) reaction time across participants, number of targets and probabilities. (**C**): Reaction time as a function of the approach direction in equiprobable (blue trace) and unequiprobable (green trace) session (**D**): Reaction time as a function of target separation computed from single-target trials across 3 participants. Error bars correspond to SE, solid lines show the polynomial regression fitting (linear in panels A and D, quadratic and cubic in panels B and C) and the colored shadow areas illustrate the confidence interval of the polynomial regression results. Target probability influences both the approach direction and the reaction time of the reaches. However, trial-by-trial analysis showed that reaction time and approach direction are not fully mediated by the target probability. Instead, reaches with longer reaction times often launch to an intermediate location between the potential goals.

By plotting RT vs. approach direction separately for the two sessions, we found that changes in RT are independent of target probability and accounted for by approach direction. Fig. 4C shows RT as a function of the initial approach direction across all participants and trials separately for the two sessions. Importantly, RT increases with reaches to intermediary location between the potential goal locations and peaks around 20*^°^* (possibly due to the biomechanical constraints of the reaching movements) regardless of the target probability (best fit cubic regression model; R-square *<* 0.95, p-value *<* 0.01 for the cubic coefficient in both sessions). To ensure that this effect was not due to some inherent constraints induced by the experimental setup - i.e., reaches launched to targets located at the center of the screen have longer RTs than reaches aimed to peripheral targets - we varied the target separation between 0.10 m to 0.20 m (which corresponds to a visual angle between 26.5 and 45 degrees) in the equiprobable session and computed the RT in the single-target trials. No significant association was found between target location and RT (linear regression model: R-square = 0.476, p-value = 0.197 of the linear coefficient), Fig. 4D. Instead, both approach direction and reaction time are driven by trial-by-trial variations in a latent variable, which we identify with decision confidence as we describe in the following sections. Note that in this analysis we used 3 individuals, who were not part of the main experiment and did not go through the training session before running the task. This could explain why RTs were slightly longer compared to single-target trials in the two main sessions.

### 2.4 Action selection, reaction time and choice confidence emerge through action competition

Our results require a decision computation that would produce joint changes in trajectory and RT as a function of trial-by-trial fluctuations in decision confidence. A recently developed theory [11, 12] predicts exactly these effects. In the theory, action decisions are made through a continuous competition of parallel prepared actions by dynamically integrated all sources of information about the quality of the alternative options. The neurodynamic implementation of this theory for a dual-target trial is presented in Fig. 5. The framework consists of a set of dynamic neural fields (DNFs), which mimic the neural processes underlying spatial sensory input, expected outcome, reach cost (i.e., effort) and reach planning [11]. Each DNF simulates the dynamic evolution of firing rate activity within a neuronal population. The functional properties of each DNF are determined by the lateral interactions within the field and the connections with other fields [18, 19].

**Figure 5.**
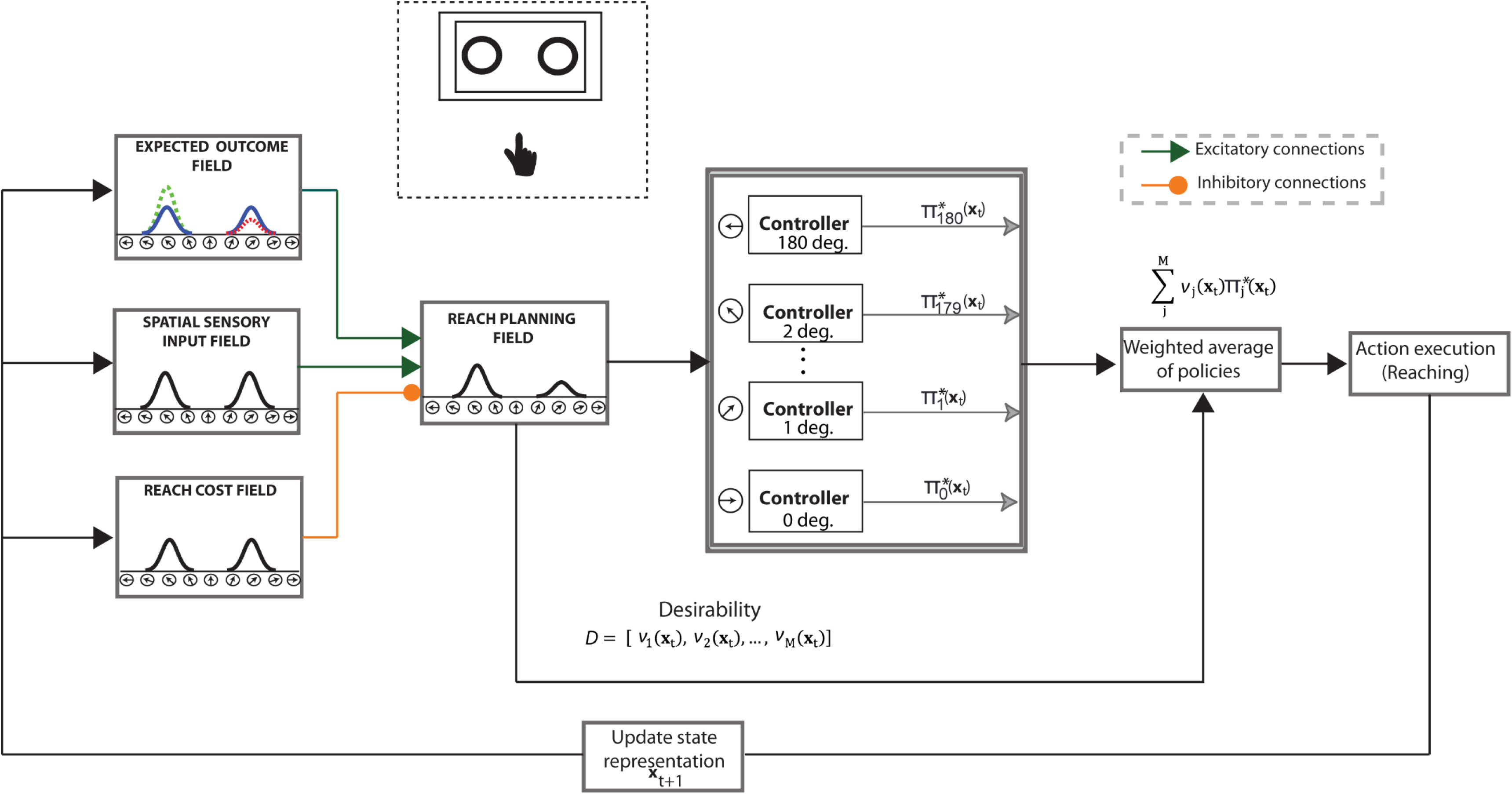
Model architecture of the “reach-before-you-know” task. The neural fields consist of 181 neurons and their spatial dimension spans the semi-circular space between 0*^°^* and 180*^°^*. Each neuron in the reach planning field is connected with a stochastic optimal control system. Once the activity of a neuron exceeds a threshold *γ*, the corresponding controller generates a sequence of reach actions towards the preferred direction of the neuron. The reach planning field receives excitatory inputs from the spatial sensory input field that encodes the angular representation of the potential targets, and the expected outcome field that encodes the expected outcome of the competing targets (blue, red and green Gaussian distributions correspond to cues with 0.5, 0.2 and 0.8 target probability, respectively). It also receives inhibitory inputs from the reach cost field that represents the effort required to move towards a particular direction. The normalized activity of the reach planning field encodes the “desirability” of the *M* available sequences of actions (i.e., neurons with activation level above the threshold *γ*) at a given time and state and acts as a weighting factor on each individual sequence of actions. Because the relative desirability is time- and state-dependent, a range of behavior from weighted averaging (i.e., spatial averaging trajectories) to winner-take-all (i.e., direct reaches to one of the cues) is generated.

The “reach planning” field employs a neuronal population code over 181 potential movement directions to plan motor actions towards these directions. It receives one-to-one excitatory inputs from the “spatial sensory input” field that encodes the angular representation of the targets and the “expected outcome” field that represents the expected outcome of aiming to a particular direction. It also receives one-to-one inhibitory inputs from the “reach cost” field that encodes the effort required to move to a particular direction. Each neuron in the reach planning field is projected to a stochastic optimal control system. Once the activity of a reach neuron *i* exceeds a threshold *γ* at the current state **x***_t_*, the corresponding controller initiates an optimal sequence of actions (i.e., policy, *π*^*∗*^) to move the “hand” towards the preferred direction of that neuron (see materials and methods section for more details). The normalized activity of the reach planning field represents the *desirability* of the motor actions, and acts as a weighting factor on them. It reflects how “desirable” it is to move to a particular direction with respect to the alternatives. Because desirability is time- and state-dependent, the weighted mixture of individual actions automatically produces a range of behavior, from direct reaching movement to weighted averaging.

Fig. 6A illustrates the activity of the planning field as a function of time for a representative dual-target trial with equiprobable targets. Initially, the field activity is in the resting state. After targets onset, two neuronal populations selective for the targets are formed and compete through mutual inhibitory interactions, while integrating information about the target certainty and action cost to bias the competition. Once the activity of one them exceeds a response threshold, the corresponding target is selected and a reaching movement is initiated. Frequently, the neuronal activity of the unselected target is not suppressed before movement onset, resulting in reaches towards intermediary locations between the targets (top inset in Fig. 6A). After the movement onset, the two neuronal ensembles retain activity and compete against each other until the goal onset.

**Figure 6.**
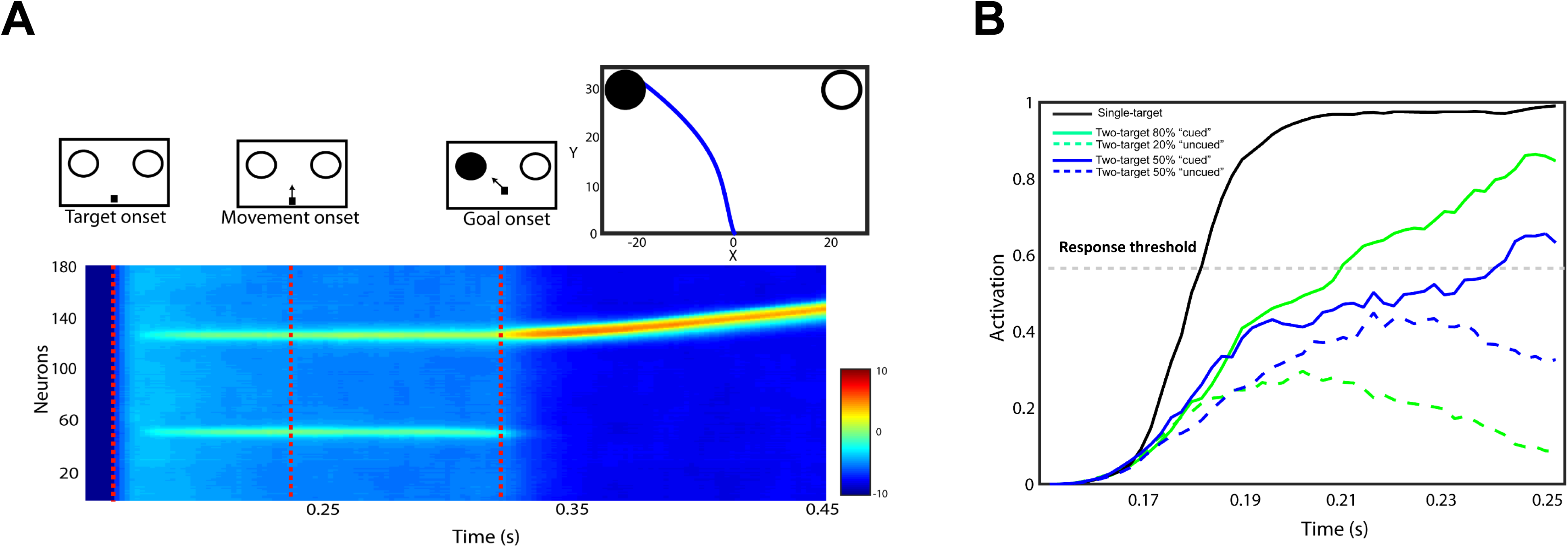
Simulated neural activity and reach behavior. (**A**): A representative example of the simulated model activity as a function of time in the reach planning field for a dual-target trial with the actual goal located in the left visual field. The red discontinuous lines indicate the target onset, the movement onset, and the goal onset. The corresponding reach trajectory is shown in the upper inset. (**B**): Simulated activity of two planning neurons centered at the location of the cued (continuous traces) and the uncued (discontinuous traces) target, from a representative single-target trial (black trace) and two dual-target trials with equal (blue traces) and unequal (green trace) probabilities. A reach movement is initiated when the activity of one of the neurons exceeds the response threshold (gray discontinuous trace). When only a single target is presented, the neuronal activity ramps up quickly to the response threshold resulting in faster reactions and direct reaches to the target. However, when two targets are simultaneously presented, the neurons compete for selection through inhibitory interactions resulting often in slower reaction times and spatially averaged movements. If one of the alternatives is assigned with higher probability, the competition is biased to the likely target leading to faster responses.

To get better insight on the model computations consider two neurons, one from each population, centered at the target locations. Fig. 6B depicts the activity of each neuron (i.e., which reflects its current desirability value) as function of time for a dual-target trial with equal (blue traces) and unequal (green traces) target probability. The neuron that exceeds the response threshold first (continuous traces) dictates the reaction time and the selected target. Intuitively, if the race between the neurons is a close call (blue traces), it means that the net evidence supporting that the selected target is more desirable than the alternative is weak and therefore individuals should be less confident about their choices. On the other hand, if the race was a landslide (green traces), it means that one alternative outperforms the other and therefore individuals should be more confident about their choice. Going back to the population analysis, the “winning” population determines the reaction time and the selected target, whereas the “losing” one contributes to the computation of the confidence that the selected option is the best current alternative. Note that in the absence of action competition (i.e., single-target trials), the activity of the neuron exceeds the response threshold faster than when two actions compete for selection (black trace). Hence, reaches have shorter RTs and aim directly to the goal location. Overall, the theory is analogous to the normative race models in perceptual decisions in which two accumulators integrate sensory evidence in favor of two alternative options [4, 20]. The accumulator that reaches its upper bound faster dictates the reaction time and the choice, whereas the losing accumulator contributes to the computation of certainty that the choice is correct.

We simulated the two equiproble and unequiprobable sessions within the computational theory, having the same fixed parameter values used to model a different reaching dataset [11]. Consistent with the human behavior, we found that target probability is correlated with the approach direction Fig. 7A (best fit linear regression model: R-square = 0.984, p-value = 0.0005 of the linear coefficient) and the RT Fig. 7B (best fit quadratic regression model: R-square = 0.984, p-value 0.008 of the quadratic coefficient). We also tested trial-by-trial association between RT and approach direction and found the same independence from target probability, Fig. 7C (best fit cubic regression model: R-square *<* 0.970, p-value *<* 0.007 for the cubic coefficient in both sessions). In particular, simulated reaches aimed towards an intermediary location between the potential targets had longer RT than reaches launched closer to one of the competing options regardless of the target probability. This is explained by the inhibitory competition between the neuronal ensembles that slows down the reach onset and leads to spatial averaging movements, if the population of the unselected action is not completely suppressed at the movement initiation. Considering that the difference between the desirability values determines the confidence of the selected action suggests that approach direction and RT are not fully coupled but there is a third variable (i.e., confidence level) that influences the association between them. That is, the longer that you wait to make an action, the less confident you are feeling about the selected action, because often the unselected one is not fully rejected. Overall, our findings provide direct evidence that action selection, reaction time and confidence that the selected action is better than the alternatives emerge through a common mechanism of desirability-driven competition between parallel prepared actions.

**Figure 7.**
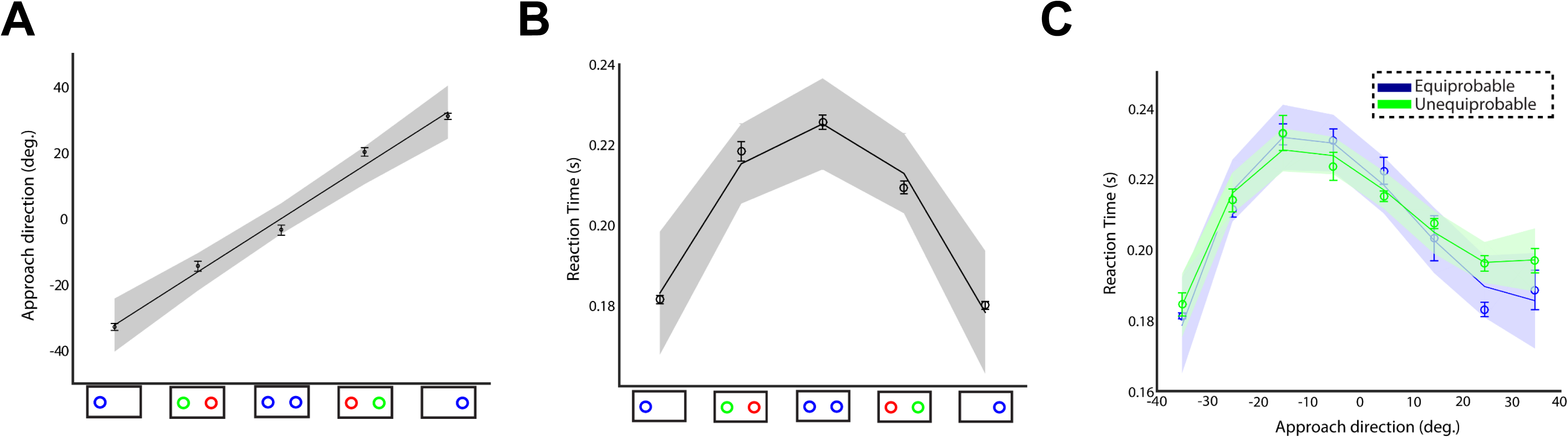
Approach direction and reaction time of the simulated reaches. (**A**): Approach direction and (**B**) reaction time of the simulated reaches across number of targets and probabilities. (**C**): Reaction time as a function of the approach direction in the simulated equiprobable (blue trace) and unequiprobable (green trace) sessions. Error bars correspond to SE, solid lines show the polynomial regression fitting (linear in panel A, quadratic in panel B and cubic in panel C) and the colored shadow areas illustrate the confidence interval of the polynomial regression results. Consistent with the human findings, the model predicts that target probability influences both the approach direction and the reaction time of the movements. However, reaction time and approach direction are not fully mediated by the target probability. Instead, the longer it takes to resolve the action competition, the more likely it is the losing population to be still active at the movement onset, resulting in spatially averaged reaches.

## 3 Discussion

### 3.1 General

Uncertainty is ubiquitous in our interactions with the external world, and decisions regularly must be made in the face of it. Even after a decision is made, there is residual uncertainty that persists in the form of subjective choice certainty, reflecting the strength of our belief that an option is *better* in the sense it is more likely correct or has a higher expected outcome than it’s alternatives. Although monitoring subjective choice certainty is crucial in guiding adaptive behavior, especially in complex and dynamic environments, decision neuroscience has focused on the primary problem of predicting decisions, largely ignoring the meta-cognitive role of monitoring decision quality. This was partly due to lack of reliable measurements to estimate confidence, especially in nonverbal animals, and also due to a paucity of theoretical proposals for how confidence emerges in decisions.

Recent efforts have been made to fill both these gaps. Perceptual decision studies in humans have simultaneously measured choices and confidence levels [4, 21], while a postdecision wager method has been introduced to measure confidence in nonverbal animals [2, 3, 22]. In addition, normative models, which include drift diffusion, evidence-accumulation, and race models [23–28], have been extended to understand how confidence emerges in perceptual decisions [4, 21]. Although parsimonious, these studies are highly restricted and limited to binary perceptual choices made solely on the basis of the accumulation of sensory evidence in static and fixed environments. In these models, confidence is construed as reflecting the effective amount of sensory evidence at decision time, which is not adequate to account for the subjective choice certainty in complex decisions. More commonly, decisions are made in dynamic and complex environments, in which the value and the availability of the options change with time and previous actions, entangling decision with action selection. Subjective confidence should reflect all the factors that affect our belief that we have made the best choice, and we need to enlarge our conception of confidence to include these factors.

In the current study, we adopted this enriched view to explore how confidence emerges in decisions requiring reaching to targets with uncertainty. Confidence was modeled as reflecting the degree of subjective belief that a potential action is more desirable than its alternatives. We designed a “reach-before-you-know” experiment in which individuals were instructed to perform rapid reaches to one or two potential targets presented simultaneously in both hemifields. To elucidate the computations underlying confidence, we modeled the task within a recently developed computational theory [11,12]. It is based on the idea that decisions are made through a continuous competition between neuronal populations that plan individual actions to the available goals, while dynamically integrating information into a common currency - named relative desirability - to bias the competition. The desirability reflects the belief about the quality of the action and acts as weighted factor on each individual action. The neuronal population that exceeds first a response threshold dictates the reaction time and the selected target. The competing population that did not exceed the threshold contributes to the computation of the confidence; the closer the “losing” population to the threshold the lower the confidence about the selected option. When the activity of the losing population is not completed suppressed, reaches are aimed towards an intermediary location between the targets. Therefore, the approach direction is an easy-to-measure proxy for choice confidence. The model predicts a direct association between target certainty with approach direction of the initial reaches and reaction time. When both targets are equally probable, the competition between the two populations is frequently a close call, which means that the net evidence supporting the selected action is weak and we should be less confident about the current best action. This results in slower reaction times and spatially averaged movements to an intermediary location between the potential goals. On the contrary, when one of the targets is assigned with higher probability, the competition is biased to the likely target. In this case the net evidence supporting the selected action is strong and therefore we should be more confidence about the current best action. This results in faster reaction times and more direct reaches to the selected target. Importantly, the model suggests that the association between reaction time and approach direction is not fully mediated by the target certainty. Instead, the longer it takes to initiate an action, the more likely it is that the losing population will still be active at the movement onset, resulting in lower confidence level about the selected option and spatially averaged movements. Hence, reaction time and approach direction are not fully mediated by the target probability, but they are influenced by the confidence about the current best option. Overall, we provide direct evidence for the first time that action selection, reaction time and choice confidence emerge through a continuous competition between parallel prepared actions.

Consistent with the model predictions, individuals adopted a spatial averaging behavior to compensate for the goal location uncertainty. Although this behavior has been reported before [7–9], the pattern of compensation is better described as buying more time for decisions. When people are uncertain about the current best option, they delay the decision both by moving towards an intermediary location between the targets and by having a longer reaction time. In contrast, when certain they initiate movement quickly and aim directly to the selected target. In line with the model predictions, trial by trial reaction time was correlated with the approach direction regardless of the target probability. Longer reaction times are often associated with weak accumulated evidence about the current best option (i.e., strong competition between the desirabilities of the actions). This might suggest that the brain learns to use decision time as a proxy for confidence judgment (see also [4, 5, 29])

### 3.2 From sensory evidence to desirability competition

Although our theory employs an “accumulator” mechanism, it is quite different from the race models. It does not assign a priori populations of neurons to alternative options; rather the alternative options emerge within a distributed neuronal population by integrating information from multiple sources. Consequently, it can handle not only binary decisions, but also decisions between multiple competing goals. It accumulates and integrates information from more than one source (e.g., sensory evidence, expected outcome, action cost, etc). Importantly, it is not limited by the serial order assumption that action planning begins only after a decisions is made. Instead, the competing options are continuously evaluated after the movement onset, whereas a decision can be changed while acting in the presence of new information. The main difference between our theory and the normative race models is that the “accumulators” compete based on the relative desirability, instead of the sensory evidence of the alternative options. Desirability provides a more general measurement to evaluate an alternative, since it includes information not only about the goal itself, but also the action required to achieve that goal. Our theory is inline with a series of neurophysiological and pharmacological intervention studies in animals reporting that areas in the posterior parietal cortex integrate value information to estimate the relative desirability of available options [30–34]. On the contrary, although neurons in specific PPC regions exhibit activity patterns that directly resemble the evidence accumulation process posited in race models [35, 36], recent studies reported that silencing these neurons does not influence the decision process [37, 38]. These findings question the role of PPC in perceptual decisions and prompt more scrutiny of the evidence-accumulation models [39].

### 3.3 Parallel versus serial hypothesis for action selection

The key point of our theory is that the brain plans multiple actions in parallel that compete for selection, and this competition continues into execution. Although a growing body of experimental studies provide evidence in favor of parallel planning of competing actions [7, 8, 13, 40–45], other studies argued against this hypothesis suggesting that decision and action are separate processes - i.e., planning and execution of action occur after a decision is made [46–51]. According to this theory, the spatial averaging behavior observed in dual-target trials does not necessarily reflect “motor averaging” - i.e., simultaneous planning of multiple competing single-target actions - but it could be equivalently interpreted as evidence of “visual averaging” across the locations of the targets - i.e., planning and execution of a single action towards a weighted averaged target location [43, 44]. The visual averaging hypothesis could explain the spatial averaging behavior and some aspects of action selection and reach timing. For instance, it could be argued that reaction time is shorter in the unequiprobable trials because individuals aim more often directly to the likely target, instead of estimating first and then moving to the weighted average location between the targets. However, the visual averaging hypothesis is insufficient to explain how choice confidence is diminished with reaction time regardless of the target probability. This effect can be modeled only within two modules that accumulate and integrate sources of information in favor of the two options and compete for selection (see an analogous case for perceptual decisions in [4]).

The action competition hypothesis is also in apparent conflict with a recent study arguing that planning and initiation of an action are mechanistically independent [52]. According to this study, reaction time does not reflect the time at which the competition between the parallel planned actions is resolved - i.e., there is no causal relationship between planning and initiation of actions. Instead, reaction time is determined by an independent initiation process. It is likely that action initiation occurs at a fixed delay after the action planning. However, this study did not account for goal location uncertainty or multiple competing goals. Instead, the individuals had to perform center-out reaches to one of eight peripheral targets arranged in a circle, and therefore they did not need to generate multiple actions that compete for selection. Overall, our findings provide further evidence in favor of the affordance competition hypothesis suggesting that the process of deliberating between different actions emerges via a continuous competition between these actions.

## 4 Materials and Methods

### 4.1 Participants

Seven right-handed (20-30 years old, 4 men and 3 women) individuals with normal or corrected-to-normal vision participated in this experiment study. The appropriate institutional review board approved the study protocol and informed consent was obtained based on the Declaration of Helsinki.

### 4.2 Experimental setup

A rough sketch of the experimental setup used in this study is shown Fig. 1. Participants were seated facing a Phantom Premium 1.5 Haptic Robot (Sensable Technologies, MA) and a computer display, aligned so that the midline of their body was in line with the center of screen and robot. The workspace of the phantom haptic robot forms a hemisphere approximately 30 cm in radius. The participants selected a comfortable position and inserted the right index finger into the endpoint of the tip of the robotic manipulandum. The distance *d*_*subject*_ from the head of the participants to the finger starting position measured along the *y* axis was about 0.30 m. This distance was slightly varied between participants, since we did not use a chin rest or any other restraining device. Hence, there was some movement of the head relative to the screen, but was minimal since the participants were instructed to remain stationary throughout the experiment. The distance from the finger starting position to the screen display *d*_*display*_ was about 0.35 m and was calibrated at the beginning of each session.

The participants were trained to perform rapid reaching movements using the robotic manipulandum. The reaching movements were performed in the horizontal plane and translated into movements of a small cursor circle (1.5 cm diameter) in the vertical plane of the computer screen - i.e., reaches towards the screen moved the cursor to the top of the screen, while left and right mapping was preserved. This experimental set up allowed for high temporal and spatial resolution of the hand and finger position as well as a mean to create haptic feedback or altered movement dynamics for future experiments. Control of the phantom robot and the experiment were implemented using the OpenHaptics drivers provided by Sensable technologies, and the Simulation Laboratory (SL) and Real-Time Control Software Package [53] as well as other custom psychophysics software. Control and recording of the phantom state were performed at 500 Hz.

### 4.3 Experimental paradigm

At the start of each trial participants were required to move the cursor to the starting position, located at the origin of our coordinate system, Fig. 2. A fixation cross was then presented at the center of the screen and the participants were instructed to fixate for a short period of time (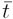 = 1500 ms, *σ*_*t*_ = 300 ms). During the final 300 ms of fixation, either a single cue was presented on the upper-left or upper-right of the screen or two cues were presented simultaneously in both sides of space. Cues were presented as unfilled circles with 3 cm in radius on a white background. After the fixation offset (go-signal) the participants had to initiate a rapid reaching movement. Once the cursor exceeded a certain trigger threshold (i.e., a virtual wall in the *x - z* plane; red discontinuous line in Fig. 2), the single cue or one of the two cues was filled-in black indicating the actual location of the goal. If the participants brought the cursor to the cued target within 1.0 s the trial was considered successful. Trials in which the participants responded before the go-signal or arrived to the cued target after the allowed movement time were aborted and were not used for further analysis. The distance between the origin and the midpoint of the two targets was *d*_*reach*_ = 0.20 m. The target separation distance - i.e., distance between the target and the midpoint - was *d*_*separation*_ = 0.15 m. The trigger threshold distance - i.e., distance of the virtual wall from the origin - was *d*_*threshold*_ = 0.05 m.

Individuals were familiarized with the task by running a set of training trials that included reaches to single and two targets. Once they felt ready and comfortable with the experimental setup, the actual experiment started. Each participant performed 3 reaching sessions (one training and two tests). The training session involved 40 trials, which were excluded from the analysis, followed by two test sessions with 80 trials each (2 *×* 80 = 160 trials). The first test session involved reaches to one (40% of the trials) and two (60% of the trials) targets. In the single-target trials, the cue was shaded blue and was presented equiprobably to the left or right visual field (top row in Fig. 2). In the two-target trials, the cues were also shaded blue and had equal probability of filling-in after the movement onset (bottom row in Fig. 2). The second test session was similar to the first one with the only difference that one of the cues was always assigned with higher probability in the two-target trials. The “likely” cue was shaded green and had 80% probability of being the correct target, while the alternative cue was shaded red and had 20% probability. The set of target configurations is illustrated in Fig. 2B. Individuals were not informed what the coloration indicates and learned the association during the experiment. Each participant performed the experiment twice with a minimum interval of 24 hours.

### 4.4 Behavioral data analysis

Cubic interpolating splines were used to smooth the reach trajectories and compute the velocity of the movements. The initial approach direction was measured from the direction of the main axis of the covariance ellipse that describes the spatial variation of the cursor from the movement initiation to the goal onset. Reaction time was defined as the time at which the reach velocity exceeded 5% of the maximum velocity.

### 4.5 Neurodynamical framework

In the current section, we briefly describe the architecture of the computational framework used to model the reaching experiment. Readers can refer to [11,12] for more details. The framework combines dynamic neural field (DNF) theory with stochastic optimal control (SOC) theory and includes circuitry for perception, expected outcome, selection bias, effort cost and decision making. Each DNF simulates the dynamic evolution of firing rate activity of a network of 181 neurons over a continuous space with local excitation and surround inhibition. The functional properties of each DNF are determined by the lateral inhibitions within the field and the connections with other fields in the architecture. The projections between the fields are topologically organized - i.e., each neuron *i* in a field drives the activation at the corresponding neuron *i* in the other field. The activity of a DNF evolves over time under the influence of external inputs, local excitation and lateral inhibition interactions as described by Eq. (1)

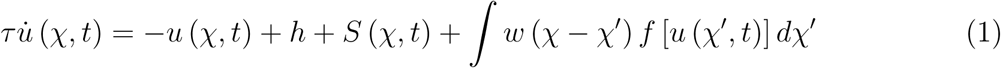

where *u* (*χ, t*) is the local activity of the DNF at the position *χ* and time *t*, and 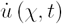 is the rate of change of the activity over time scaled by a time constant *τ*. If there is no external input *S*(*χ, t*), the field converges over time to the resting state *H* from the current level of activation. The interactions between the simulated neurons in the DNF are given via the kernel function *w* (*χ - χ^’^*), which consists of both local excitatory and inhibitory components, Eq. (2).

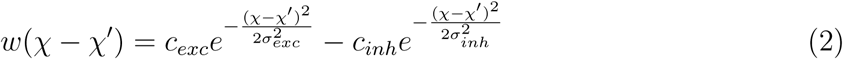

where *c*_*exc*_, *c*_*inh*_, *σ*_*exc*_, *σ*_*inh*_ describe the amplitude and the width of the excitatory and the inhibitory components, respectively.

We convolved the kernel function with a sigmoidal transformation of the field so that neurons with activity above a threshold participate in the intrafield interactions, Eq. (3).

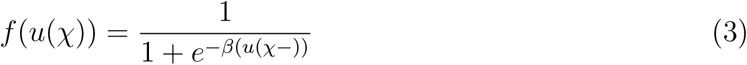

The architectural organization of the framework is shown in Fig. 5. The “spatial sensory input” field encodes the angular representation of the competing goals in an ego-centric reference framework. The expected outcome for reaching to a particular direction centered on the hand position is encoded by the “expected outcome” field (see [11] for more details). In trials with equiprobable targets, the neuronal activity of the populations selective for these targets is about the same (blue Guassian distributions). However, in trials in which one of the targets is more likely than the alternative, the activity of the neuronal population selective for the “green” cue is higher than the activity of the populations which is tuned to the “red” cue. The outputs of these two fields send excitatory projections (green arrows) to the “reach planning” field in a topological manner. The “reach cost” field encodes the effort cost required to move to a particular direction at any given time and state. The output of this field sends inhibitory projections (orange arrow) to the reach planning field to penalize high-effort actions. The activity of the reach planning field at a given state **x***_t_* is sum of the outputs of the fields encoding the location of the target **u***_loc_*, the expected outcome **u***_outcome_* and the estimated reach cost **u***_cost_*, corrupted by additive noise *ξ* which follows a Normal distribution.

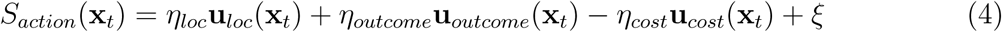

where *η*_*loc*_, *η*_*outcome*_ and *η*_*cost*_ scale the influence of the spatial sensory input field, the expected outcome field and the reach cost field, respectively, to the activity of action planning field. The values of the model parameters are given in the *S*_1_, *S*_2_ and *S*_3_ of the supporting information in [11]. The normalized activity of the action planning field describes the “relative desirability” of each policy *π*_*i*_ - i.e., it reflects how “desirable” it is to move towards a particular direction *φ*_*i*_ with respect to the alternative options.

Each neuron in the reach planning field is linked with a stochastic optimal controller. Once the activity of a neuron *i* exceeds a threshold *γ*, the controller *i* is triggered and generates an optimal policy 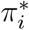 - i.e., sequence of actions towards the preferred direction of the neuron *i* - which is given by minimizing the following cost function:

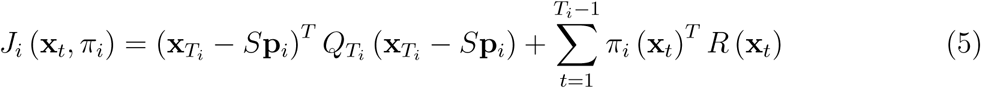

where the policy *π*_*i*_ (**x***_t_*) is a sequence of actions from *t* = 1 to *t* = *T*_*i*_ to move towards the direction *ϕ*_*i*_; *T*_*i*_ is the time required to arrive at the position **p***i*; **p***i* is the goal-position at the end of the movement and is given as **p***i* = [*r* cos(*ϕ*_*i*_), *r* sin(*ϕ*_*i*_)], where *r* is the distance between the current location of the hand and the location of the cue which is tuned by the neuron *i*. Additionally, **x***_T_i* is the state vector at the end of the movement, whereas the matrix *S* picks out the actual position of the hand and the goal-position **p***i* at the end of the movement from the state vector. Finally, *Q*_*T*_*i* and *R* define the precision- and the control-dependent cost, respectively. For more details about the optimal control model used in the framework see the supporting information in [11, 12].

The first term of Eq.(5) describes the current goal of the controller - i.e., move the hand at a distance *r* from the current location, towards the preferred direction *ϕ*_*i*_ of the neuron *i*. The second term describes the cost required for executing the policy *π*_*i*_ (**x***_t_*). Let’s now assume that *M* neurons are active at a given time *t* (i.e., the activity of *M* neurons is above the threshold *γ*). The framework computes and executes a weighted average of the individual policies 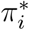 to move the hand from the current state **x***_t_* to a new one, Eq. (6).

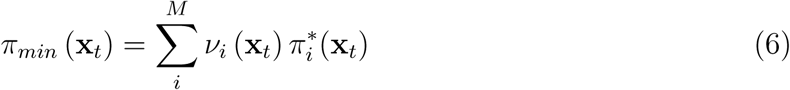

where *v*_*i*_(**x***_t_*) is the normalized activity of the neuron *i* (i.e., the relative desirability value) at the state **x***_t_*. Because the desirability is time- and state-dependent, the weighted mixture of the individual policies produces a range of behavior, from winner-take-all (i.e, direct reaching to a target) to spatial averaging.

To handle contingencies, such as perturbations (e.g., changes on the number of targets, target probabilities, expected rewards, etc) and effects of noise, the framework implements a widely used technique in stochastic optimal control known as “receding horizon” [54,55]. In particular, the framework executes only the initial portion from the sequence of actions for a short period of time *k* (*k* = 10 in our study) and then recomputes the individual optimal policies 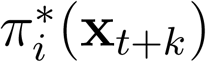 from time *t* + *k* to *t* + *k* + *T*_*i*_ and remixes them. This approach continues until the hand arrives to one of the targets.

## References

1. Foote AL and Crystal JD. Metacognition in the rat. Curr Biol., 17(6):551–555, 2007.

2. Hampton RR. Rhesus monkeys know when they remember. Proc Natl Acad Sci U S A., 98(9):5359–5362, 2001.

3. Kepecs A, Uchida N, Zariwala HA, and Mainen ZF. Neural correlates, computation and behavioural impact of decision confidence. Nature, 455(7210):227–231, 2008.

4. Kiani R, Corthell L, and Shadlen MN. Choice certainty is informed by both evidence and decision time. Neuron, 84(6):1329–1342, 2014.

5. Fetsch CR, Kiani R, Newsome WT, and Shadlen MN. Effects of cortical micros-timulation on confidence in a perceptual decision. Neuron., 83(4):797–804, 2014.

6. De Martino B, Fleming SM, Garrett N, and Dolan RJ. Confidence in value-based choice. Nat Neurosci., 16(1):105–110, 2013.

7. Chapman CS, Gallivan JP, Wood DK, Milne JL, Culham JC, and Goodale MA. Reaching for the unknown: Multiple target encoding and real-time decision-making in a rapid reach task. Cognition, 116(2):168–176, 2010.

8. Gallivan JP, Chapman CS, Wood DK, Milne JL, Ansari D, Culham JC, and Goodale MA. One to four, and nothing more: Nonconscious parallel individuation of objects during action planning. Psychol Sci., 22(6):803–811, 2011.

9. Hudson TE, Maloney LT, and Landy MS. Movement planning with probabilistic target information. J neurophysiol., 98(5):3034–3046, 2007.

10. Gallivan JP and Chapman CS. Three-dimensional reach trajectories as a probe of real-time decision-making between multiple competing targets. Front Neurosci., 8(215), 2014.

11. Christopoulos V, Bonaiuto J, and Andersen RA. A biologically plausible computational theory for value integration and action selection in decisions with competing alternatives. PLoS Comput Biol., 11(3), 2015.

12. Christopoulos V and Schrater PR. Dynamic integration of value information into a common probability currency as a theory for flexible decision making. PLoS Comput Biol., 11(9), 2015.

13. Cisek P. Cortical mechanisms of action selection: the affordance competition hypothesis. Philos Trans R Soc Lond B Biol Sci., 362(1485):1585–1599, 2007.

14. Cisek P and Kalaska JF. Neural mechanisms for interacting with a world full of action choices. Annu Rev Neurosci., 33(1):269–298, 2010.

15. Beck JM, Ma WJ, Kiani R, Hanks T, Churchland AK, Roitman J, Shadlen MN, Latham PE, and Pouget A. Probabilistic population codes for bayesian decision making. Neuron, 60(6):1142–1152, 2008.

16. Pleskac TJ and Busemeyer JR. Two-stage dynamic signal detection: a theory of choice, decision time, and confidence. Psychol Rev., 117(3):864–901, 2010.

17. Stewart BM, Gallivan JP, Baugh LA, and Flanagan JR. Motor, not visual, encoding of potential reach targets. Curr Biol., 24(19):953–954, 2014.

18. Erlhagen W and Schöner G. Dynamic field theory of movement preparation. Psychol Rev., 109(3):545–572, 2002.

19. Schöner G. Cambridge Handbook of Computational Cognitive Modeling, chapter Dynamical systems approaches to cognition, pages 101–126. Cambridge University Press, 2008.

20. Vickers D and Packer J. Effects of alternating set for speed or accuracy on response time, accuracy and confidence in a unidimensional discrimination task. Acta Psychol., 50(2):179–197, 1982.

21. van den Berg R, Anandalingam K, Zylberberg A, Kiani R, Shadlen MN, and Wolpert DM. A common mechanism underlies changes of mind about decisions and confidence. Elife, 5(e12192), 2016.

22. Kiani R and Shadlen MN. Representation of confidence associated with a decision by neurons in the parietal cortex. Science., 324(5928):759–764, 2009.

23. Vickers D and Smith P. Accumulator and random-walk models of psychophysical discrimination: a counter-evaluation. Perception, 14(4):471–497, 1985.

24. Usher M and McClelland JL. The time course of perceptual choice: the leaky, competing accumulator model. Psychol Rev., 108(3):550–592, 2001.

25. Gold JI and Shadlen MN. Banburismus and the brain: decoding the relationship between sensory stimuli, decisions, and reward. Neuron, 36(2):299–308, 2002.

26. Mazurek ME, Roitman JD, Ditterich J, and Shadlen MN. A role for neural integrators in perceptual decision making. Cereb Cortex., 13(11):1257–1269, 2003.

27. Krajbich I and Rangel A. Multialternative drift-diffusion model predicts the relationship between visual fixations and choice in value-based decisions. Proc Natl Acad Sci U S A., 108(33):13852–13857, 2011.

28. Towal BR, Mormann MM, and Koch C. Simultaneous modeling of visual saliency and value computation improves predictions of economic choice. Proc Natl Acad Sci U S A, 110(40):3858–3867, 2013.

29. Hanks TD, Mazurek ME, Kiani R, Hopp E, and Shadlen MN. Elapsed decision time affects the weighting of prior probability in a perceptual decision task. J Neurosci., 31(17):6339–6352, 2011.

30. Sugrue LP, Corrado GS, and Newsome WT. Matching behavior and the representation of value in the parietal cortex. Science, 304(5678):1782–1787, 2004.

31. Dorris MC and Glimcher PW. Activity in posterior parietal cortex is correlated with the relative subjective desirability of action. Neuron, 44(2):365–378, 2004.

32. Wardak C, Olivier E, and Duhamel JR. Saccadic target selection de?cits after lateral intraparietal area inactivation in monkeys. J Neurosci., 22(22):9877–9884, 2002.

33. Wilke M, Kagan I, and Andersen RA. Functional imaging reveals rapid reorganization of cortical activity after parietal inactivation in monkeys. Proc Natl Acad Sci U S A, 109(21):8274–8279, 2012.

34. Christopoulos VN, Bonaiuto J, Kagan I, and Andersen RA. Inactivation of parietal reach region affects reaching but not saccade choices in internally guided decisions. J Neurosci., 35(33):11719–11728, 2015.

35. Shadlen MN and Newsome WT. Neural basis of a perceptual decision in the parietal cortex (area lip) of the rhesus monkey. J. Neurophysiol., 86(4):1916–1936, 2001.

36. Gold JI and Shadlen MN. Neural computations that underlie decisions about sensory stimuli. Trends Cogn Sci., 51(1):10–16, 2001.

37. Erlich JC, Brunton BW, Duan CA, Hanks TD, and Brody CD. Distinct effects of prefrontal and parietal cortex inactivations on an accumulation of evidence task in the rat. eLife, 10.7554, 2015.

38. Katz LN, Yates JL, Pillow JW, and Huk AC. Dissociated functional signi?cance of decision-related activity in the primate dorsal stream. Nature, 535(7611):285–288, 2016.

39. Pesaran B and Freedman DJ. Where are perceptual decisions made in the brain? Trends Neurosci., 39(10):642–644, 2016.

40. Basso M and Wurtz R. Modulation of neuronal activity in superior colliculus by changes in target probability. J Neurosci., 18(18):7519–7534, 1998.

41. Glimcher PW, Dorris MC, and Bayer HM. Physiological utility theory and the neuroeconomics of choice. Games Econ Behav., 52(2):213–256, 2005.

42. Rangel A and Hare T. Neural computations associated with goal-directed choice. Curr Opin Neurobiol., 20(2):262–270, 2010.

43. Cisek P. Making decisions through a distributed consensus. Curr Opin Neurobiol., 22(6):927–936, 2012.

44. Gallivan JP, Barton KS, Chapman CS, Wolpert DM, and Flanagan JR. Action plan co-optimization reveals the parallel encoding of competing reach movements. Nat Commun., 6(7428), 2015.

45. Gallivan JP, Logan L, Wolpert DM, and Flanagan JR. Parallel specification of competing sensorimotor control policies for alternative action options. Nat Neurosci., 19(2):320–326, 2016.

46. Friedman M. Essays in Positive Economics. Chicago University Press, Chicago, IL, 1953.

47. Tversky A and Kahneman D. The framing of decisions and the psychology of choice. Science, 211(4481):453–458, 1981.

48. Fodor JA. Modularity of Mind: An Essay on Faculty Psychology. MIT Press, Cambridge, MA, 1983.

49. Pylyshyn ZW. Computation and Cognition: Toward a Foundation for Cognitive Science. The MIT Press, Cambridge, MA, 1984.

50. Padoa-Schioppa C and Assad JA. Neurons in orbitofrontal cortex encode economic value. Nature, 44(7090):223–226, 2006.

51. Padoa-Schioppa C. Neurobiology of economic choice: a good-based nodel. Annu Rev Neurosci., 34:333–359, 2011.

52. Haith AM, Pakpoor J, and Krakauer JW. Independence of movement preparation and movement initiation. J Neurosci., 36(10):3007–3015, 2016.

53. Stefan Schaal. The *sl* simulation and real-time control software package. Technical report, University of Southern California, http://wwwclmc.usc.edu/publications/S/schaal-TRSL.pdf, 2009.

54. Mayne DQ, Rawlings JB, Rao CV, and Scokaert PM. Constrained model predictive control: Stability and optimality. Automatica, 36(6):789–814, 2000.

55. Goodwin GC, Seron MM, and de Dona JA. Constrained control and estimation: an optimisation approach. Springer, London, UK, 2005.

